# The Plankton Imager (Pi-10): An imaging instrument for automated and continuous zooplankton data collection

**DOI:** 10.1101/2025.01.27.635045

**Authors:** James Scott, Sophie G. Pitois, Robert Blackwell, Hayden Close, Phil Culverhouse, Julian Tilbury

## Abstract

The Pi-10 is the latest iteration of the Plankton Imager: a high-speed colour line-scan camera that images particles in a flow-through system. The Pi-10 is a cost-effective, easy to install and low maintenance automated instrument that can be used on any platform with access to water and power supply. We tested the Pi-10 on the research vessel Cefas Endeavour, connected to a continuous water supply pumping water at 34 L min-1. The instrument collected images of particles, within the size range of 180 µm - 3.5 cm, automatically and continuously, alongside other vessel operations, in all weathers over a period of 18 days. The Pi-10 successfully captured and saved up 5000 images per minute, translating into a 46 GB of digital storage per day. When particle density exceeded 147 per litre, the instrument stopped saving all images, while still recording the number of particles that passed through the system. This is akin to subsampling, with more sub-sampling required in areas or times of high particle density (e.g at the time of spring plankton bloom or in turbid waters). The Pi-10 collects high volumes of data in a continuous manner, thus providing unprecedented fine spatial data. The high frequency nature of the instrument opens the door to new areas of research. These include, in particular, the observation of fine scale processes, the move towards real-time sampling, and the increased capability to build a digital twin of the oceans. As technologies continue evolving the Pi-10 performance will increase, being able to collect and save more and more images.

## 1 Introduction

Zooplankton are a critical component of pelagic ecosystems. They occupy a central position in the food web, often controlling smaller organisms by grazing and providing food for many important larval and adult fish as well as seabirds (Lauria et al., 2013; Pitois et al., 2012). Their short life cycle renders zooplankton sensitive to environmental changes (Edwards and Richardson, 2004; Harris and Edwards, 2014; Serranito et al., 2016), which makes them ideal candidates as indicators of climate change. Decades of laboratory and field investigations have shown major impacts of changing oceans on zooplankton physiology, community composition, distribution and the resulting influence on both biogeochemistry and productivity of the oceans (Ratnarajah et al., 2023).

Traditionally, the collection of zooplankton samples is performed by deploying nets and subsequent analysis by an expert taxonomist using light microscopy. This is a labour intensive, time-consuming, and expensive process (Benfield et al., 2007; Wiebe and Benfield, 2003). The increasing demand for zooplankton data combined with ever reducing budgets for monitoring have driven the development of new tools and techniques for the sampling and analysis of this key ecosystem component (Pitois et al., 2023). A suite of instruments revolves around in situ imaging of zooplankton combined with machine learning algorithms to automatically classify the collected images into taxonomic groups. Such examples include: the Underwater Vision Profiler (UVP, Barth and Stone, 2022; Drago et al., 2022; Picheral et al., 2010), In Situ Ichthyoplankton Imaging System (ISIIS, Cowen and Guigand, 2008; Panaïotis et al., 2022), Video Plankton Recorder (VPR, Davis et al., 2005), and Lightframe On-sight Key species Investigation system (LOKI, Schulz et al., 2010).

Here, we describe the Plankton Imager (Pi-10), a high-speed, colour, line-scan camera that images all particles continuously in a through-flow sampling system. The Pi-10 is unique as it is connected to a continuous water supply (from a ship or any other platform) and can work automatically 24/7 alongside other operations in all weathers. One major advantage is that there is no deployment and thus no extra effort needed, in relation to labour and ship time, to collect zooplankton information as part of a survey. All images collected by the Pi-10 are saved to disk to be classified later either manually by an expert taxonomist or via a machine learning algorithm selected by the user to answer their needs.

The Pi-10 is the product of 11 years of development from 2011 to 2022. There have been several iterations of the instrument with regular upgrades to hardware and software from its initial configuration (Figure 1). The first iteration was the Line-scanning Zooplankton Analyser (LiZA) (Culverhouse et al., 2016; Culverhouse et al., 2015). The LiZA was developed to capture monochrome images of mesozooplankton and provide taxonomic information via its associated Plankton Image Analyser (PIA) computer vision system, so that data were immediately available without the need for significant amounts of post-cruise processing and analysis, thus reducing the burden on the human taxonomist. Subsequently, the instrument was renamed ‘Plankton Imager Analyser’ (PIA) in 2016, combining the image capture and analysis steps together. The main evolution was the inclusion of a colour camera system. The PIA was first tested on the RV Cefas Endeavour in autumn of 2016 and its performance compared and evaluated against two other methods: the Continuous Automated Litter and Plankton Sampler (CALPS) which sampled the same water feed as the PIA and the traditional vertical deployment of a ring net. The results showed good agreement in the spatial distribution of abundances of zooplankton across the three systems but some differences in catchability between taxonomic entities were noted (Pitois et al., 2018).

**Figure 1.**
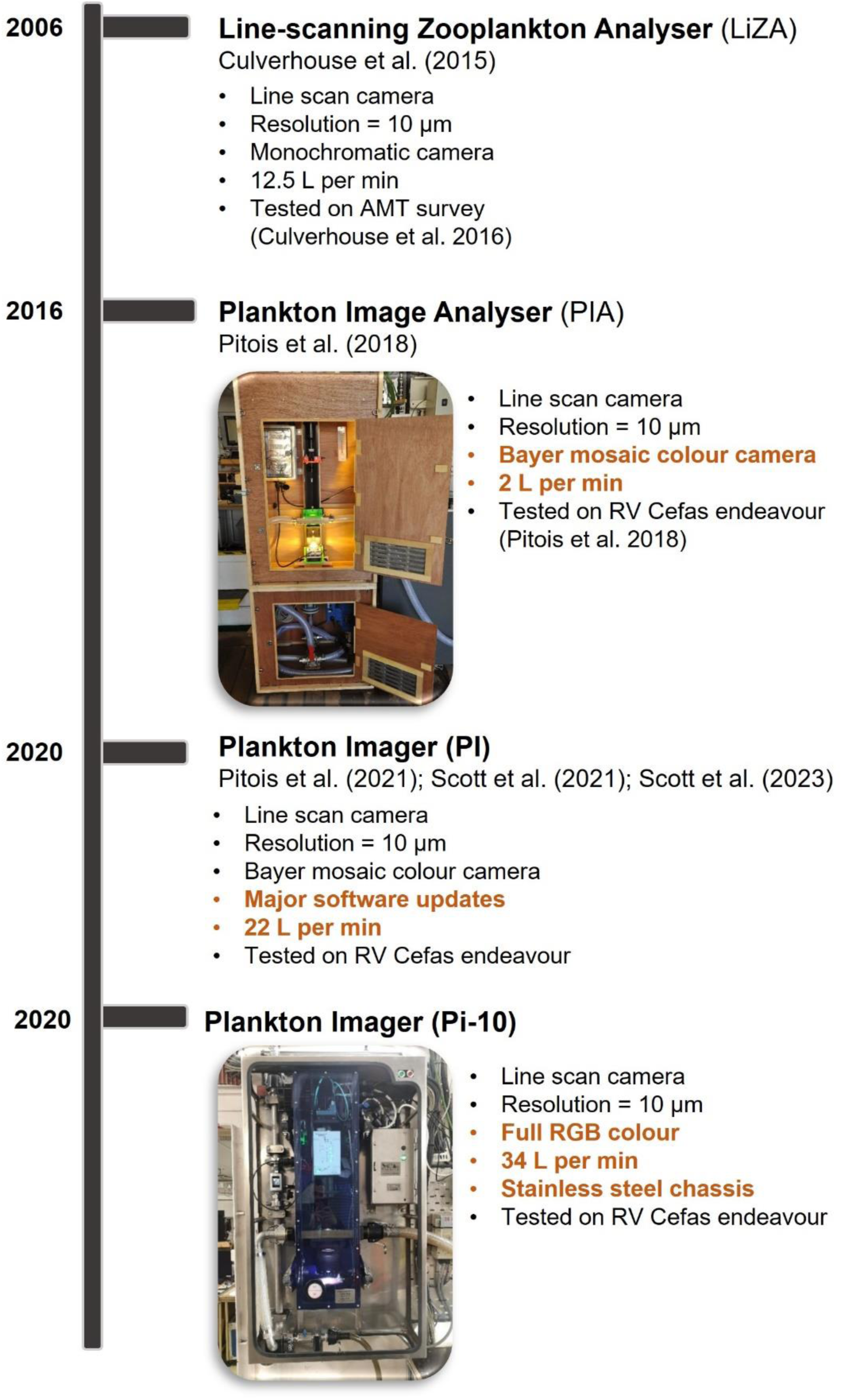
Timeline with the four major iterations of the Plankton Imager; including photographs of the instrument installed on a ship and key hardware configurations. Modifications from the previous iteration are highlighted in bold orange characters.

These were in part due to blurring of specimen features resulting from limited PIA optical depth of field, thus rendering positive classification of many images impossible. Improvements to the camera system were made to increase the depth of field and thus improve image focus and led to the next development of the renamed Plankton Imager (PI). Testing of the PI led to studies on application to food web size-based indicators (Pitois et al., 2021) and changes in mesozooplankton community structures (Scott et al., 2021). Results from these studies demonstrated: (1) the PI’s value in collecting routine in-situ sizes over many individual plankters, allowing for a more realistic picture of size distribution within a population, and resulting in more robust size-based metrics; And (2) the PI’s ability to describe temporal changes in the mesozooplankton community and capture the seasonality of individual taxa. This was found to be in line with outputs from sampling programmes in the same area. Finally, following replacement of the camera and lens, the commercially available, finished Plankton Imager was completed at the end of 2022 and named Pi-10, reflecting its 10 µm pixel resolution. Overall, the Plankton Imager has evolved from being a camera, lens and flow cell assembled together (PIA, PI), to being a standalone instrument with a microprocessor-based controller with internal climate control to maintain a constant temperature around the optics and reduce the humidity within the enclosure.

Whilst the use of images collected by the Plankton Imager to answer ecological questions has been demonstrated in previous publications (Pitois et al., 2018, 2021; Scott et al., 2021, 2023), the focus of the present study is on the technical performance of the instrument itself. Specifically, this paper aims to provide a comprehensive description of the Pi-10 instrument and its functioning, as an aid to potential users. We also provide the result of a test performed to evaluate the performance of the system, in relation to the number and rate of images captured and saved as well as their sizes.

Those results are used to discuss the instrument strengths and limitations, the potential applications for the monitoring and study of zooplankton, and thus to help the potential user to make an informed decision on how to best use the instrument.

## 2 Materials and Equipment

The Plankton Imager Pi-10 consists of two mains parts, a camera unit (Figure 2) and a controlling computer (PiPC) (Figure 3). The camera unit comprises a line scan camera, a flow cell, an LED and a control box all housed in a sealed, waterproof (ip67 rating) acrylic housing (ii in Figure 2). The sealed unit also prevents dust fouling the glass of the flow cell, the camera lens and the LED. The acrylic box is mounted on rubber mounts to reduce vibrations and subsequent loss of image quality which are within a permanently mounted stainless-steel frame (i in Figure 2). This allows the instrument to be easily removed from the ship as needed. The stainless-steel frame also houses the required plumbing, flowmeter and a power supply unit which are connected to the camera unit when in use.

**Figure 2.**
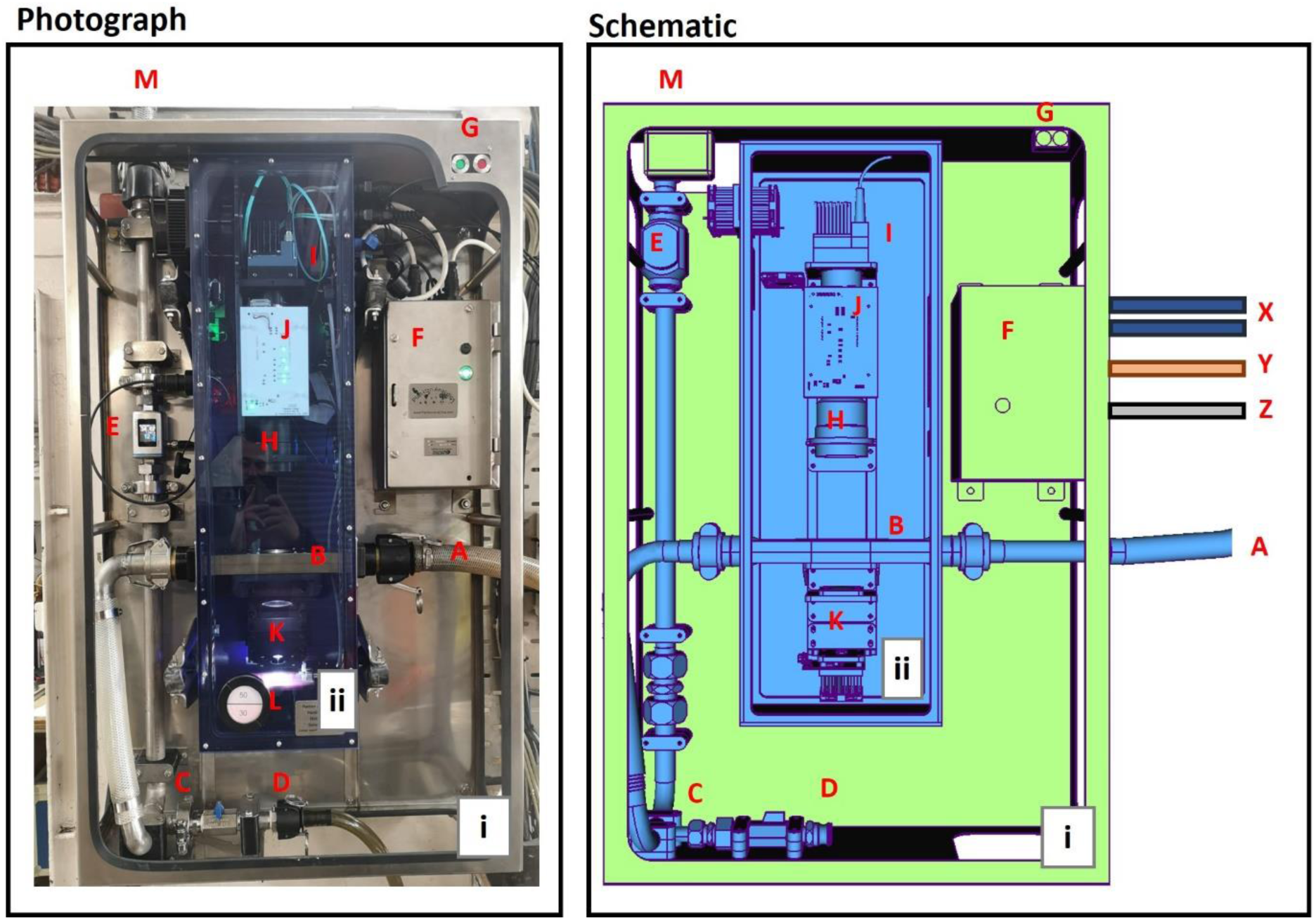
Photograph of the Plankton Imager (Pi-10) camera unit on the RV Cefas Endeavour (left) and associated schematic. The camera unit (ii) is removable and mounted on a permanent steel chassis (i). A: Water inlet to acrylic housing from the ship’s continuous flow; B: flow cell; C: Drain valve; D: Drain outlet; E: Flowmeter; F: Power supply unit; G: LED displays indicating normal operation of instrument (green) or fault; H camera lens; I: Line scan camera; J: control board shows component instrument status; K: LED light source for illumination of flow cell; L: Humidity meter; M: Water outlet; X: Fibre optic cables; Y: USB controlling cable to the controlling computer (PiPC); Z: Power lead.

**Figure 3.**
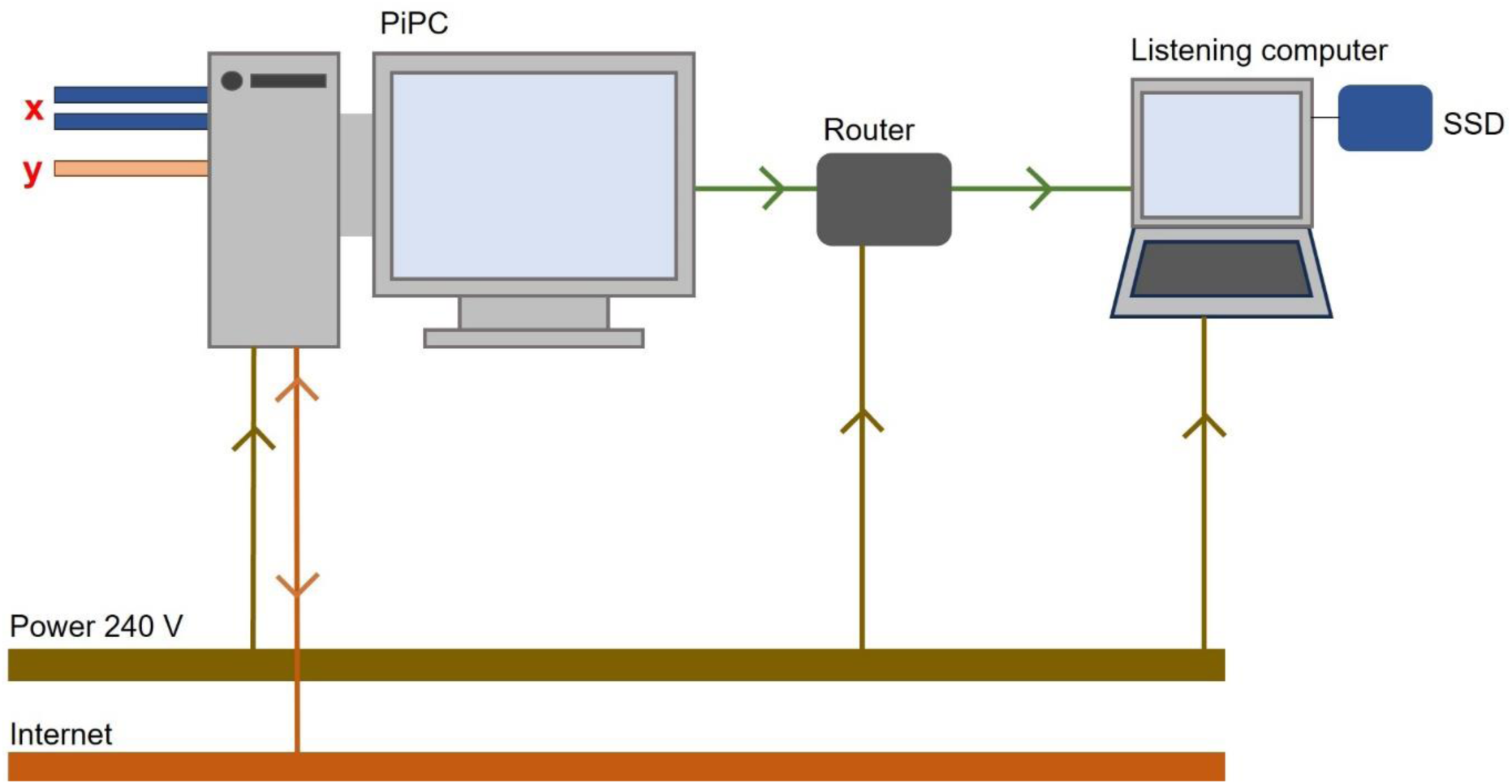
Cable schematic for the Pi-10 controlling computer (PiPC). X: Fibre optic cables and Y: USB cable controlling cable from the camera unit (Figure 2). Mains 240 V power cables (brown lines) supply all computers and respective peripherals. Internet (orange lines), provided by the ship’s network is connected only to the PiPC for remote access. Extracted images are passed over a local area network (green lines) from the PiPC using a router. Multiple listening computers can connect to the router to listen to the image feed from the network.

Two fibre optic cables and a USB cable (X, Y, in Figure 2) connect the camera unit to the PiPC (X, Y, in Figure 3). On the RV Cefas Endeavour, the PiPC is behind this bulkhead in a weatherproof dry lab. The PiPC has bespoke software which is designed to: (1) control the physical environment of the camera and monitor competent status (e.g., LED temperature, flow rate) via the USB cable; (2) control the camera unit and receive the camera feed via the fibre optic cables; and (3) extract regions of interest (ROIs) from the camera feed and save them as images. These images are the primary output of the Plankton Imager.

Each component is explained in more detail below.

### 2.1 System Design – Imaging System Hardware

#### 2.1.1 Camera and lens

The camera (H in Figures 2, A in Figure 4) is a high speed Dalsa (Teledyne Dalsa, Waterloo, ON, Canada) DALSA LINEA ML-FC-08K10T line scan camera set at 4096 pixels, each 10 µm2, in full RGB colour at 24-bit resolution. The scan rate is set to 100k lines per second and data is transferred from the camera to the computer interface via a Cameralink CLHS twin optical fibres (as described above) at a data rate of 1.144 Gigabytes per second. The lens (B in Figure 4) is a telecentric macro lens of magnification 1:1 custom-made by VS-Technology (Tokyo 106-0041, Japan). The lens is fixed in aperture and focus so there is no need for any manual setup prior to use. The optical path is shown on Figure 4. Calibration beads were used to confirm the magnification factor in the size range 0.71-0.80 mm grey neutral density (for suspension in water) polythene calibration beads (Cospheric, USA).

**Figure 4.**
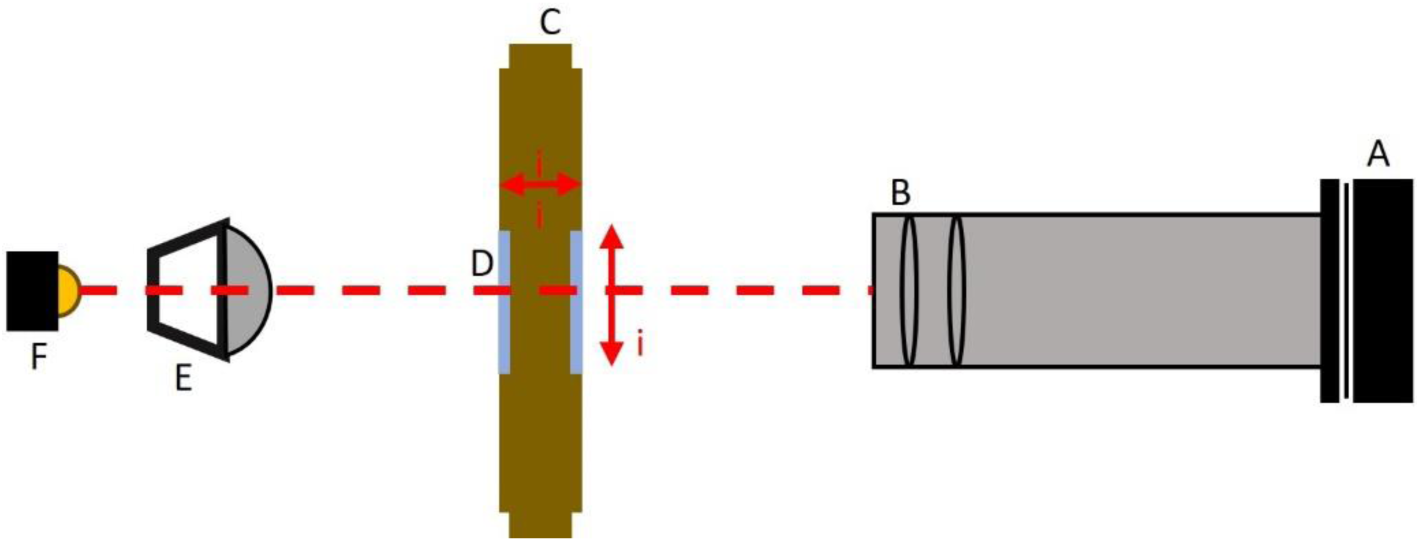
Optical path (red dashed line): A: Dalsa line scan camera; B: Telecentric macro lens; C: Brass flow cell, dimensions = 40.8 mm (i) x 13.8 mm (ii); D: Glass panel in flow cell to allow light through; E: Achromatic condenser; F – 1 mm LED.

#### 2.1.2 Flow-cell

The flow cell (B in Figure 2, C in Figure 4, Figure 5) is engineered to present a smooth transition from the water feed 32mm internal diameter hose connection to a rectangular cross-section at the imaging point in the centre of the cell. It has a 13.8 mm depth of field by 40.8 mm width, supporting 34 L.min-1 water throughput. Sapphire optical windows are used to withstand abrasion from particles suspended in the water stream. The patent (GB104989.5) details the smooth transition from circular to rectangular cross-section in the flow cell. This is designed to minimise water pressure changes, as these can induce gas bubbles formation, and standing waves, which in turn can modify the velocity of particles within the water stream.

**Figure 5.**
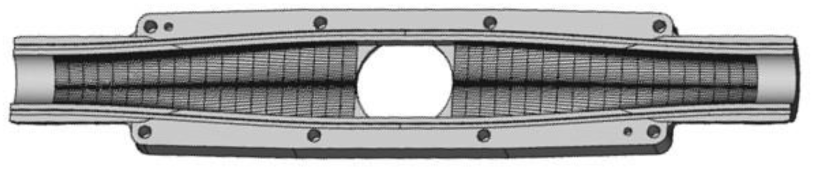
Brass flow cell computer aided design. The schematic demonstrates the transition from a smooth to angular cross section. Patent: GB2104989.5 entitled: “A flow cell and use thereof” in the name of: Plankton Analytics Limited.

#### 2.1.3 Illumination

A LED (K in Figure 2, F in Figure 4) provides white light at a colour temperature of approximately 6700k at 9400 lux into the camera sensor, which is under computer control and set upon startup Figure 6C). This can be toggled between low and high mode from the PiPC software to avoid bright light when cleaning the flow-cell. A 1 mm pinhole in line with the LED improves the light beam collimation.

**Figure 6.**
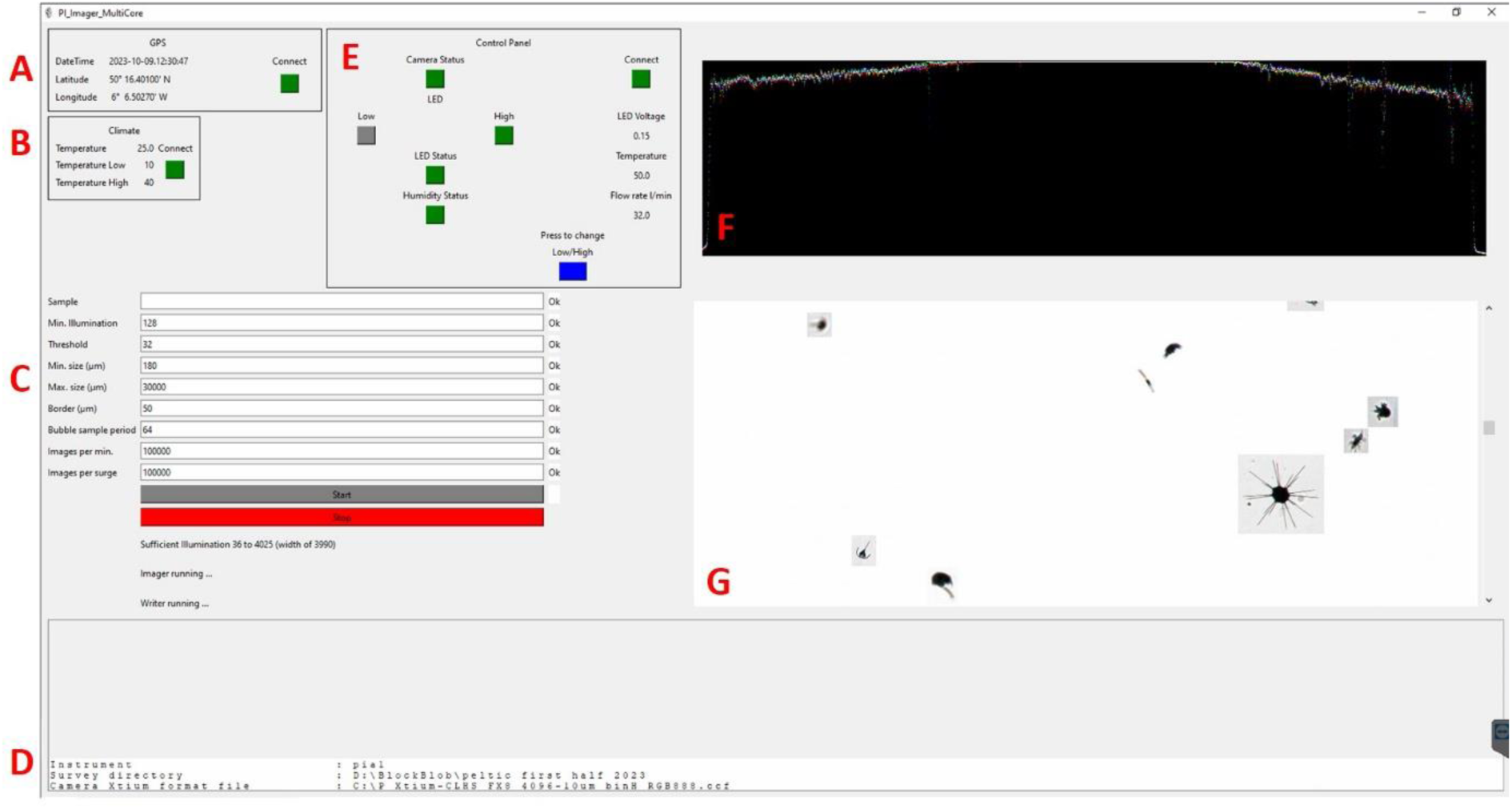
Plankton Imager controlling software (PI_Imager.exe) graphical user interface (GUI). The software controls the camera, the image parameters and displays a real-time feed of plankton images. A: time and GPS derived from the ships GPS feed via serial line; B: Temperature within the control box; C: Adjustable parameters, including illumination, threshold, minimum and maximum size of images recorded, maximum number of images recorded per minute and per surge; D: Trouble shooting window; E: A mirror of the control panel board in the acrylic box (J in Figure 2); F: RGB histogram of captured images; G: preview window for images captured by the camera and received by the PiPC in real time.

#### 2.1.4 Flowmeter

A PicoMag DN25 Electromagnetic flowmeter (E in Figure 2) is used to monitor the flow rate. The range is between 0 - 100 L m^-1^. The water flow rate is set manually by adjusting an inlet or outlet valve in line with the pumped water source. The flow rate is reported to the PiPC and displayed on the PI_Imager software GUI (E in Figure 6).

#### 2.1.5 Pi-10 Control box

The Pi-10 has a 32-bit microcontroller within its enclosure. This includes a forward-facing LED panel that indicates the status of components (J in Figure 2), and interfaces to the PiPC via USB. The PiPC receives status data from the controller every second, including the LED brightness, its operating temperature, a water leak detection sensor output, and the flow rate from the flowmeter (E in Figure 6). A USB hub also allows the PiPC to communicate with a Peltier climate controller subsystem.

Two status LEDs reflect the operational status of the Pi-10 on the stainless-steel frame (G in Figure 2). A green light indicates normal operation. Flashing green reflects a non-optimum flow rate (outside of ± 10%). Red indicates system failure, which is either an overheated illuminator, in which case the power level is automatically reduced, (flashing red) or a water leak (constant red).

#### 2.1.6 Controlling PiPC

Several computer configurations have been explored to ensure the maximum number of images can be captured and broadcast. The current system uses an AMD Threadripper with 24 cores with 32 GB of DDR3 RAM and a 512 SSD drive running Windows 10 professional. The instrument uses parallelism to achieve its throughput. Nine threads on the 24 core PiPC are dedicated to image capture, segmentation and other related activities. These are allocated automatically to cores by the computer in its OS kernel. The operating system cannot keep up with the image file creation rate of more than 100 per second. Instead, images are broadcast over UDP (User Datagram Protocol) across a local area network. “Listening” computers can connect to the LAN and receive the image stream (Figure 3). This allows the data to be accessed in real time.

All functions of the Pi-10 camera can be controlled by the PiPC via the Pi-10 software and user interface (Figure 6). The PiPC must be connected to a serial port feed for GPS streaming from the ship’s navigation system, when used onboard a vessel. Specific parts of the software are detailed below.

### 2.2 System Design – Software

#### 2.2.1 Image Acquisition Software

The Pi-10 software runs under Windows 10 operating system. The GUI allows for setting illuminator brightness, monitoring illuminator temperature, water flow rate and water leak detection within the ip67 enclosure (E in Figure 6). Some parameters can be set by the user. These include illumination (this can be used to exclude areas of the flow cell that are too dark, typically the edges), threshold (determines which pixels are dark enough to be considered to belong to particle), minimum and maximum size of particles recorded, maximum number of images recorded per minute (this limits the data rate) and images per surge (this gives the instrument a buffer capacity if a dense patch of particles is encountered) (C in Figure 6). These parameters are saved to a text file at the very beginning and very end of a run. A run is a period of time where the instrument is running continuously. This means that anyone using the data can easily see the parameters used. When image collection is initiated, the software performs background removal and particle detection. Once detected, TIFF images of the particles are broadcast on the local area network and optionally saved to local disk drive (Figure 3). Every image is dated, time and GPS stamped. A text field can be added which will form part of the TIFF metadata, for example a user might want to tag each image with survey name, location or month. This is achieved using the “Sample” option on the GUI (C in Figure 6).

To allow for post automated processing of the collected images post collection (for e.g. their identification), it is necessary to add a border around the image. The software detects plankton as shadows on a light background (i.e. if a pixel is darker than the threshold set by the user, it is deemed a shadow pixel), but plankton may have appendages that are fainter than the threshold, and this border helps to expand the image to include any missed appendage. The size of this border has been consistently set at 50 µm since its first use, but the user can customise the size of the border surrounding the image (C in Figure 6). The ROI extraction protocol adds this border and then rounds the image height and width in pixels up to the nearest factor of 8. For example, a particle of 200 µm width (i.e 20 pixels) will be added a 5-pixels border, bringing the image width to 30 pixels. This number is then rounded up the next multiple of 8 or 32 pixels. Note, when the user enters the size range they are entering the minimum size of the particle, not the image size.

As images start to be received by the PiPC, the first image of every second appears in real time on the preview window (G in Figure 6), and a RGB histogram of these images is also displayed (F in Figure 6). Variations in the histograms usually indicate dirt or marks on the flow cell or that the flow cell needs cleaning from fouling. Dirt on the flow cell might impinge image quality by obscuring passing particles. Error (if any) handling is detailed in the troubleshooting window of the GUI (D in Figure 6).

#### 2.2.2 System Throughput

The Pi-10 can operate at a range of water flow rates up to a maximum of 34 L^-1^. It is recommended running the instrument at this maximum rate, which samples two cubic metres of water per hour continuously at a resolution of 10 µm across the entire depth of field. It is our experience that this volume of water can carry from less than 1,000 particles in oceanic water to more than 200,000 particles per minute in estuarine waters. In some situations of high densities of particles in the water, the PiPC cannot keep up with the high number of images captured by the camera. In other words, when the data collection rate becomes faster than the processing rate, the system quickly clogs up with images which are then unsaved and ultimately lost (see “hits” and “misses” section below). To mitigate this and limit the operating system’s burden in handling very high numbers of newly generated small files, images are saved in 10-minute folders (this period cannot be refined by the user). The maximum number of images for a 10-minute bin can be controlled by the “image per min” and “images per surge” options on the PiPC GUI, explained below (C in Figure 6). Different hardware configurations will affect the maximum number of images that can be collected in a given time period. The following should be considered when adjusting these parameters:

– Total available data storage for the survey (i.e., the amount of space on a given hard drive to store the collected images).

– Disk write speed capacity (speed at which a disk can write images, for example, the performance of a SSD type disk would be expected to be better than that of a HDD type disk).

– Area of work (e.g. estuaries are likely to present with a high density of particulate matter in contrast to offshore areas).

– Timing of the work (Spring, mainly at the spring bloom time will present with high quantities of phytoplankton particles, in contrast to Winter).

– Desired particle size range, small particles being present in higher densities than larger particles.

For our previous work using the Plankton Imager for the ecological studies (Autumn in the Celtic Sea) referred to in the introduction, a minimum particle size was set to 180 µm. This ensured a manageable data collection rate (Pitois et al. 2018, 2021; Scott et al., 2021, 2023).

#### 2.2.3 ‘Hits’ and ‘misses’

Due to the potentially vast volume of images, the PiPC software records ‘hits’ and ‘misses’ for particles within the specified size range. Hits are the number of images successfully written to disk. This is controlled by the “images per min” and “images per surge” parameters and the actual number of particles passing through the flow cell (i.e. the particle density in the sampled water).

Misses are the number of images the camera captured but the PiPC software could not write to disk, due to reasons explained above in the “System throughput” section. Numbers of “hits” and “misses” per minute are recorded in a text file within each 10-minute bin, and the “hits:misses” ratio is used to estimate particle densities in the sampled water, similarly to the procedure applied following sub-sampling a physical sample. For example, if a ten-minute bin had 1,000 hits, of which 40 images were classified as copepods and 10,000 misses, one would assume the same relative abundance of copepods within the missed images (i.e. 40 copepods per 1,000 images), assuming a random sampling regime and applying standard sub-sampling methods.

#### 2.2.4 Image broadcasting

A local area network (LAN) is used to connect ‘listening’ computers to the controlling PiPC via a router. The UDP is used to broadcast images over the LAN via Ethernet from the PiPC. In theory, these can be received through an Ethernet port anywhere on the ship’s network so the device can be monitored anywhere on the vessel. Any machine (running any operating system), and any number of machines can listen and receive the broadcast TIFF image files when connected to the same LAN (Figure 3). The benefit of using this system is that more than one ‘listening’ device can collect images and secondly, the Pi-10 functionality can be checked anywhere on board. The hits and misses text files are also broadcast over LAN.

#### 2.2.5 Image storage

Prior to 2023, post survey images were backed up on large external hard drives as previously described. More recently images have been saved on either Microsoft Azure data disk packs and backed up on large external HDDs / SSDs. Using the Azure disk packs allows for easy movement of the images to cloud storage. The Azure disk packs are posted back to the local Azure data center and uploaded to the server directly. This also allows for using Azure cloud compute for rapid analysis of the images. Data may also be stored on privately owned storage drives and later uploaded to the cloud.

## 3 In-situ assessment of instrument

The Pi-10 was tested aboard the RV Cefas Endeavour from the 11th to 28th of November 2023 around the southwestern UK coast (Figure 7). The instrument was connected to the ship water supply, pumping water from 4m depth at 34 l/min flow rate, and worked continuously during the survey, apart from some pausing to allow for regular maintenance (e.g. cleaning the flow cell). Cleaning can take up to 20 minutes to complete, as the water flow must be turned off, the flow cell drained, cleaned with a bottle brush and a mixture of water, detergent and vinegar and then flow restored.

**Figure 7.**
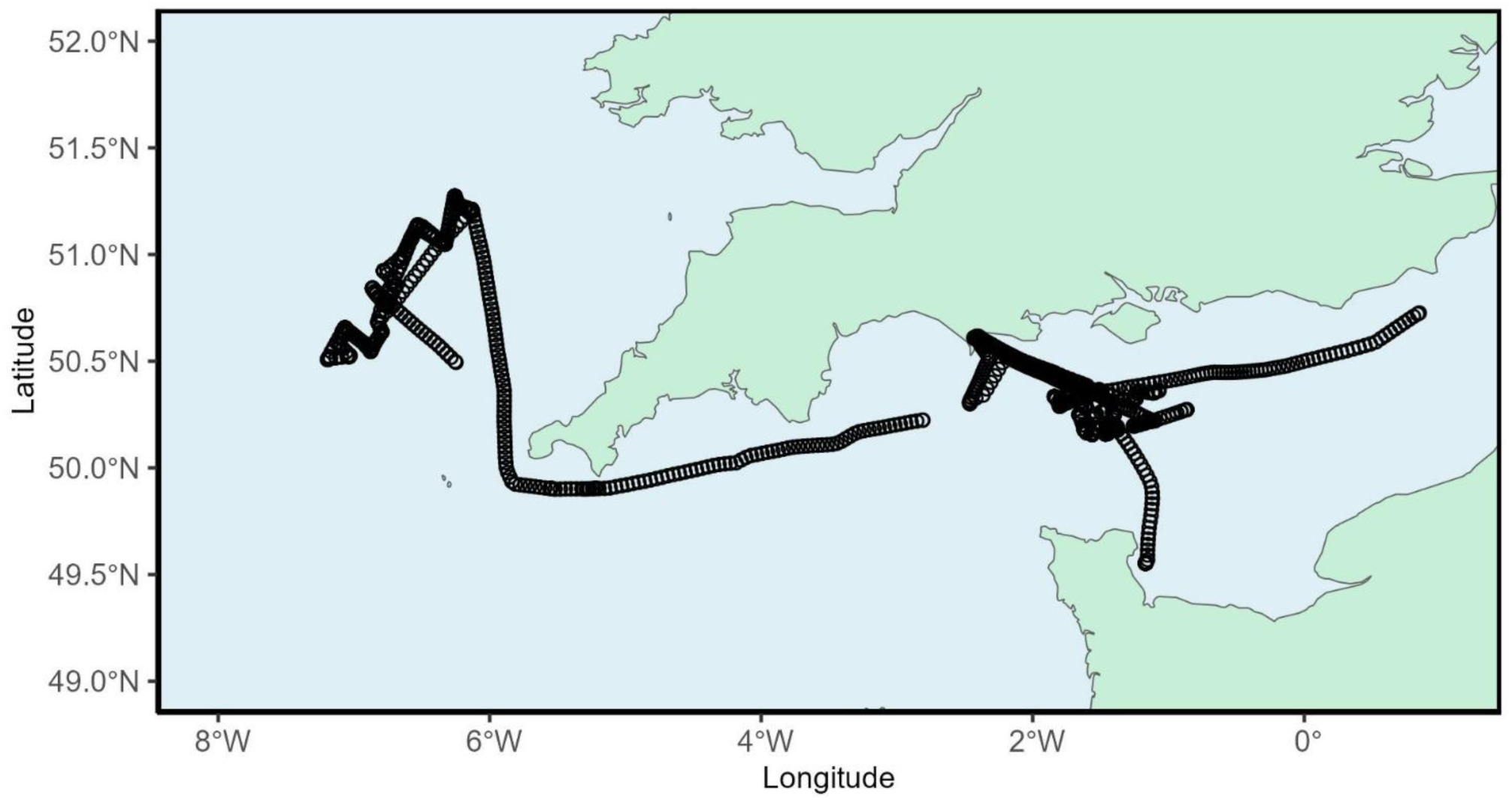
Study area around the Southwest coast of the UK. Each black circle represents ten minutes of data collection.

A size range of 180 µm to 35,000 µm was used, to ensure the capture of the entire mesozooplankton component. Images were saved to a local NTFS formatted solid state drive (SSD) disk into 10-min bins as described above. Analysis was performed on those images using a combination of Python and R.

The aim was to evaluate the total numbers and rates of images captured and successfully saved (i.e. ‘hits’ and ‘misses’) to establish the baseline performance of the instrument. An additional aim was to evaluate the dimensions of each image (height and width) and their associated size to assess the storage requirements of the collected data. To balance processing time and data requirements, analyses were focused on every one in every five 10-min bins (420 out of 2116 available directories), and one in every five images within each bin. To explore spatial patterns of hits, misses and image dimension, data were extracted into 0.2-degree grid cells.

## 4 Results

### 4.1 Image capture

Over the entire survey, the Pi-10 ran for approximately 353 hours, resulting in 2,116 ten-minute bins. A total of 70,273,294 images (See Figure 8 for examples of images collected) was successfully captured (“hits”) and 13,714,839 images could not be saved (”misses”). Therefore, the Pi-10 missed an average of 1 in 5 images overall. The mean numbers of hits and misses per bin was 33,258 and 6,490 respectively, equating to an average image capture rate of 55 per second.

**Figure 8.**
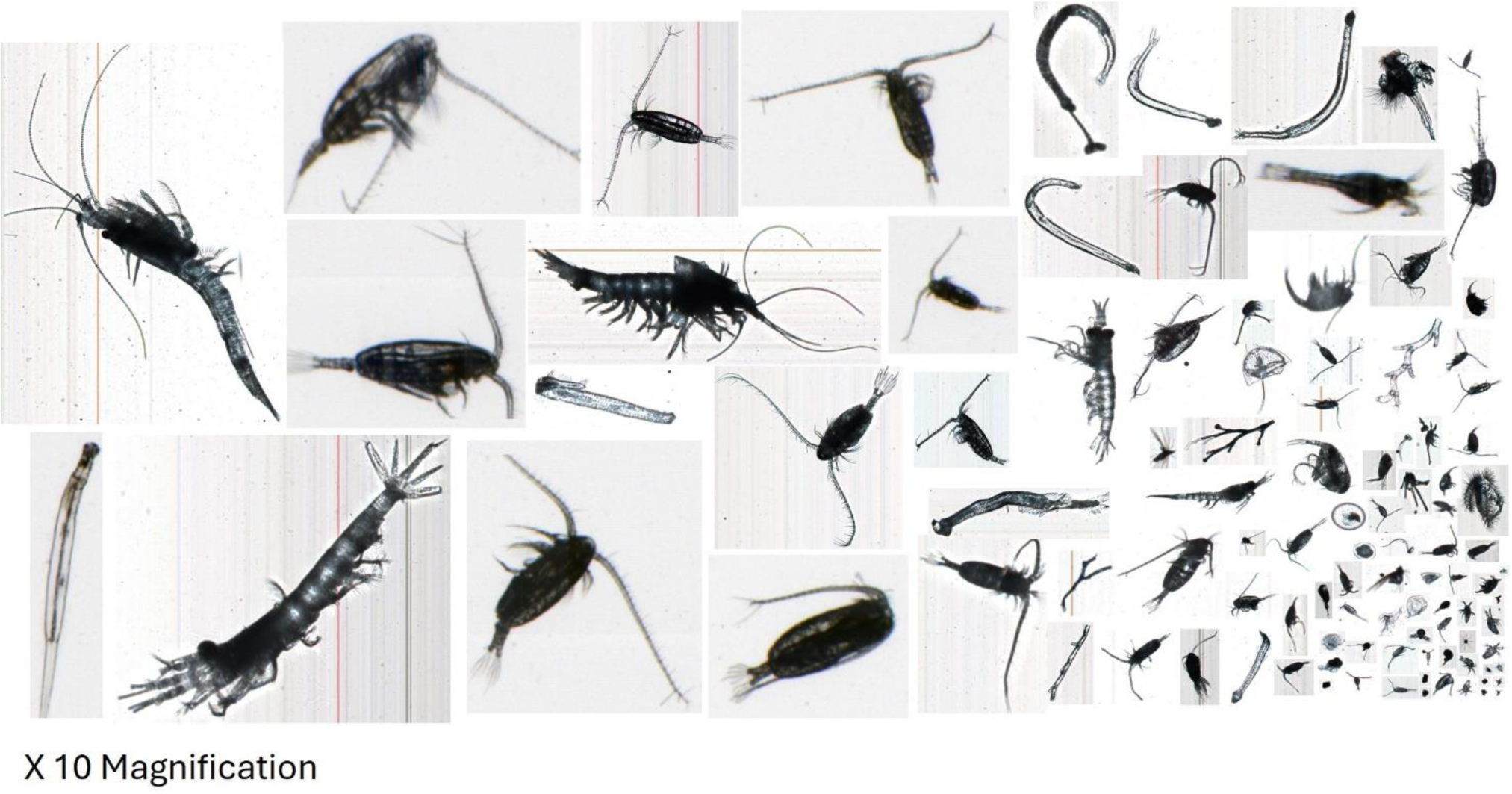
Examples of images collected by the plankton imager. Images are magnified 10 times and show zooplankton arranged from largest (left) to smallest (bottom right).

The maximum recorded number of hits in one ten-minute bin was 381,005 equating to a capture rate of 635 images a second, whilst the most misses in a single bin was 3,968,710. There were large variations between days in terms of hits and misses (Figure 9). On the 15/11/2023, data collection was not possible due to all operations on the ship being stopped. The five days from the 10/11/2023 to the 14/11/2023 had consistently below 5,000 hits per min. Following this for 5 more days to the 21/11/2023 the image capture rate increased to between 5,000 and 10,000 images per minute (Figure 9), the highest level being seen on 16/11/23. During the final week of the survey, excluding the very last day, the capture rate fell to below 2,500 images per min. Throughout the survey, misses were only recorded on 4 out of 18 days. On these four days, the capture rate exceeds 5,000 images per minute suggesting that the maximum capacity of the system sits around 5,000 images per minute. Above this rate, images are not saved. In other words, when the particle density in the water is above 147 L-1, the instrument cannot store all the images taken.

**Figure 9.**
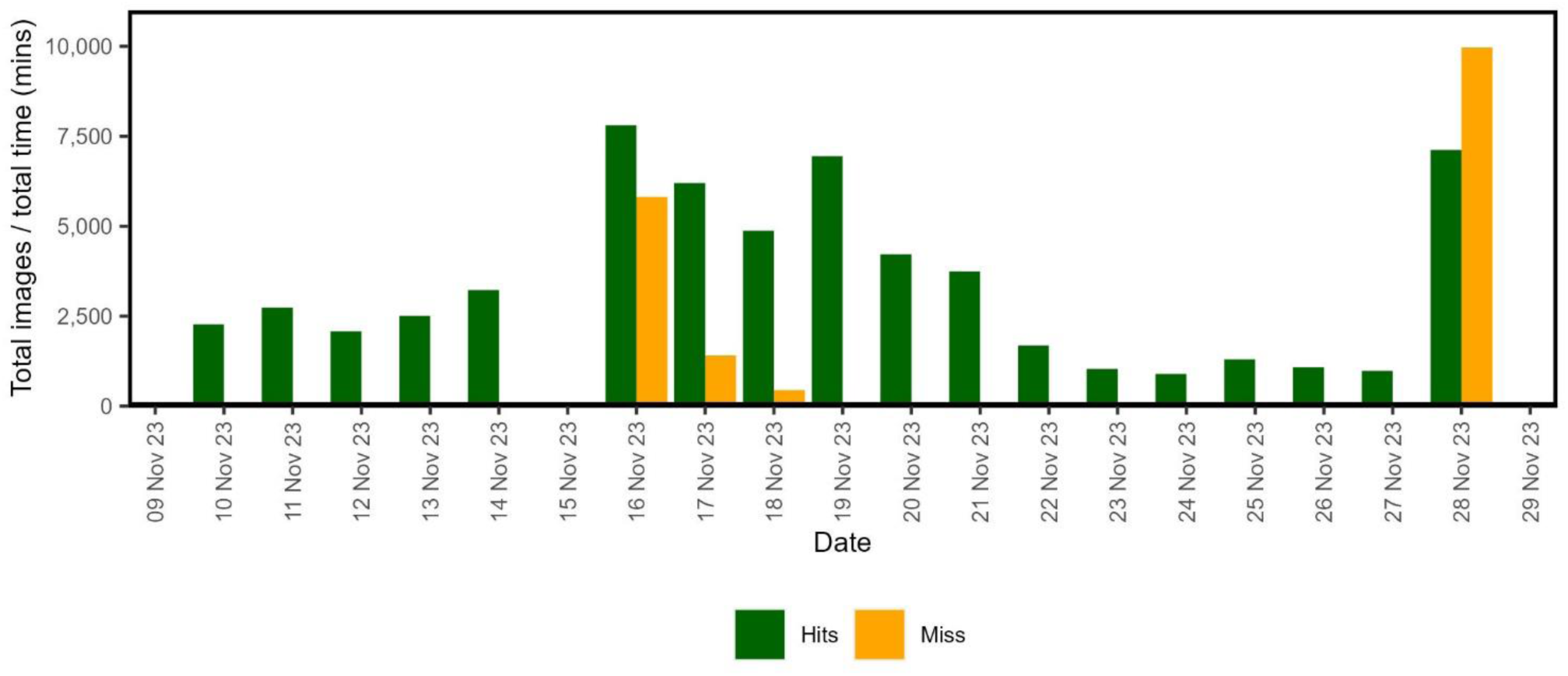
Average number of hits and misses per minute for each day when the Pi-10 was in operation calculated as total numbers within a 24h period divided by the number of minutes of operation within that period. On the 15/11/2023 no data was collected as all ship operations were stopped.

Mapping of hits and misses divided by the total time spent in each 0.2° grid cell, in general showed little variation across the study area with some notable exceptions (Figure 10). In the majority of cells, all images were saved, except for three clustered areas of high misses, one in the English Channel, south of the UK coast, another two smaller ones in the Celtic Sea, west of the UK coast (Figure 10B). Within these clusters, neighbouring cells share high numbers (> 5,000) of misses per minute. Clustered areas of misses coincide with cells with the highest numbers of hits (Figure 10A). Outside of these (i.e. when all images of particles are successfully saved), the number of hits and associated image capture rate remained consistently below 5,000 hits per minute. This confirms the above result on the system’s capacity to save no more than 5,000 images per minute. There was no correlation between the total number of images missed per bin and the mean size of either height or width, suggesting that the system does not discriminate across images when these cannot be saved.

**Figure 10.**
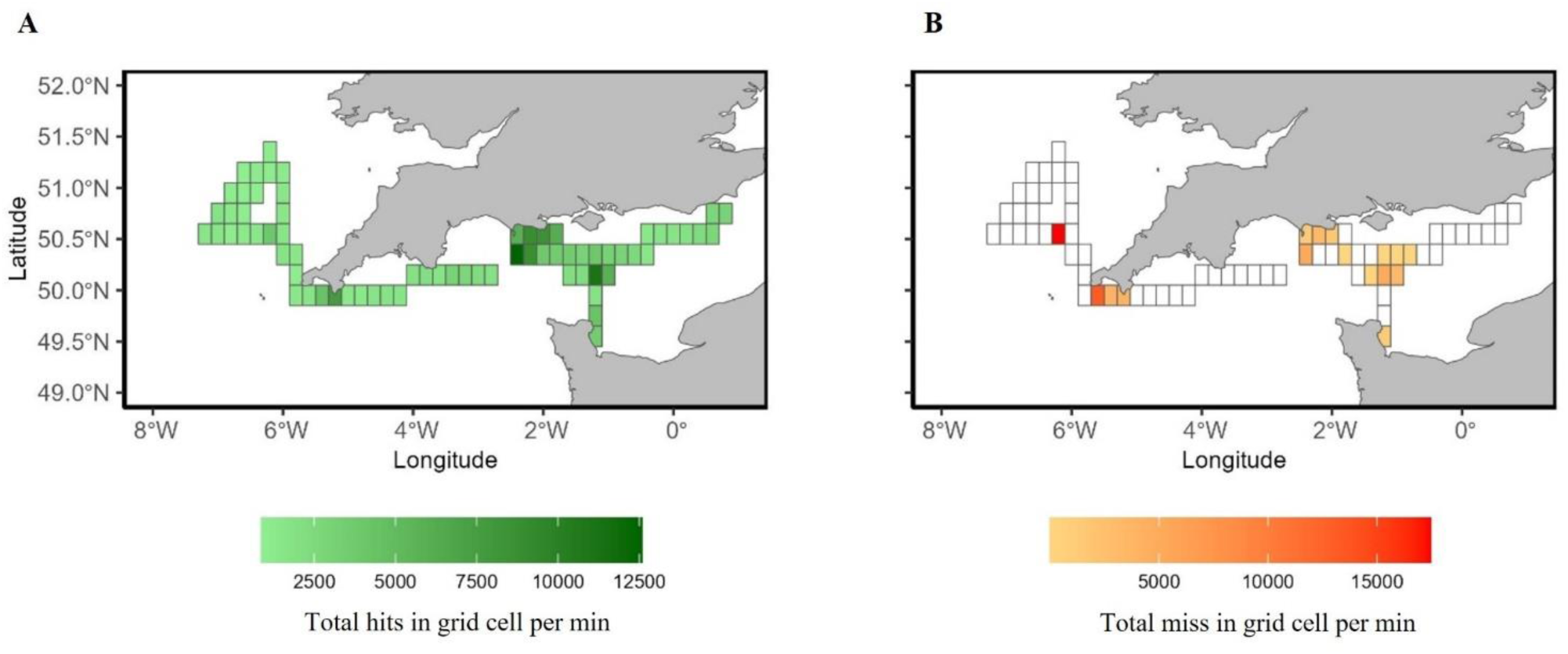
Spatial distribution of average numbers of hits (A) and misses (B) per minute. Average values were calculated as total numbers within a cell divided by the number of minutes of operation within that cell.

### 4.2 Size of images and storage requirements

The mean image height was 49.57 pixels (496 µm) and the mean image width was 47.34 pixels (473 µm). The size distribution of the collected images is skewed towards the smallest ones, with many small images and few larger ones (Figure 11). As images height or width reached 160 pixels or above, their numbers fell below 100 images in each category. These are not shown in Figure 11 to ensure clarity on the figure. Towards the very largest dimensions (> 2500 pixels) there were often only 1 or 2 occurrences.

**Figure 11.**
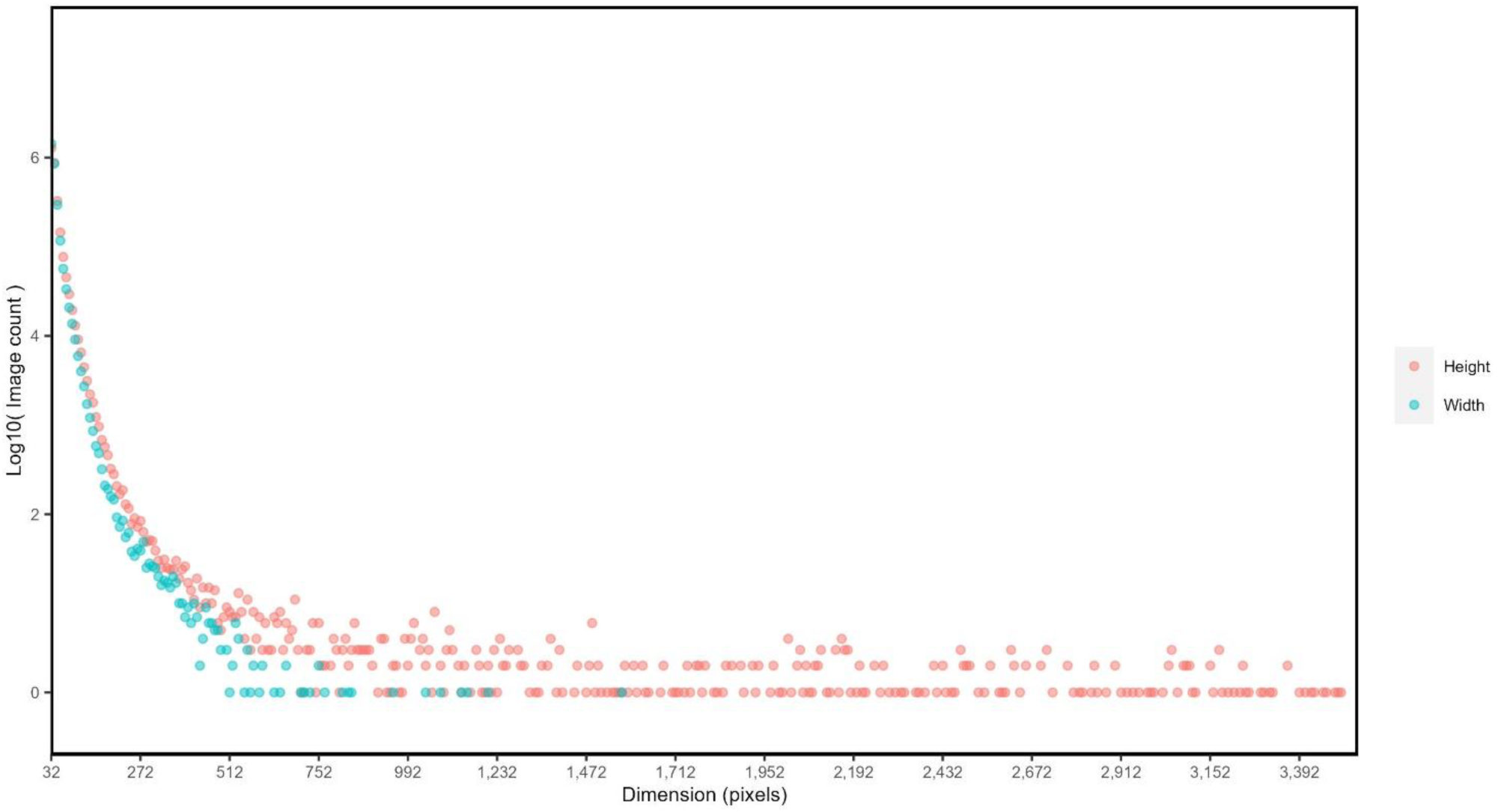
Numbers of images within 8-pixels bins for height (pink) and width (blue), from a total of 2,845,113 images collected and post-subsampling. 10 pixels = 1 µm.

Image size in terms of digital storage follows a predictable analogue with the image dimensions. The vast majority (2,346,950) of images were smaller than 0.0064 MB (Figure 12). There were only 115 large images (i.e. > 1 MB), amounting to 0.01% of the total number of images.. Assuming a mean number of hits per 10-min bin of 33,258 (or 3,3326 hits per minute) and a mean image size of 0.0064 MB, this equates to a data rate of approximately 20 MB per minute or 28,736 MB a day (28.73 GB). Therefore, a 2 TB SSD will have the capacity to store over 300,000,000 images and should last for about 70 days at sea under similar conditions to those we operated in here. There is however some variation between actual file size and required size on disk which is dependent on the disk characteristics such as format or firmware overheads (e.g. disk type, operating system used) which must be considered.

**Figure 12.**
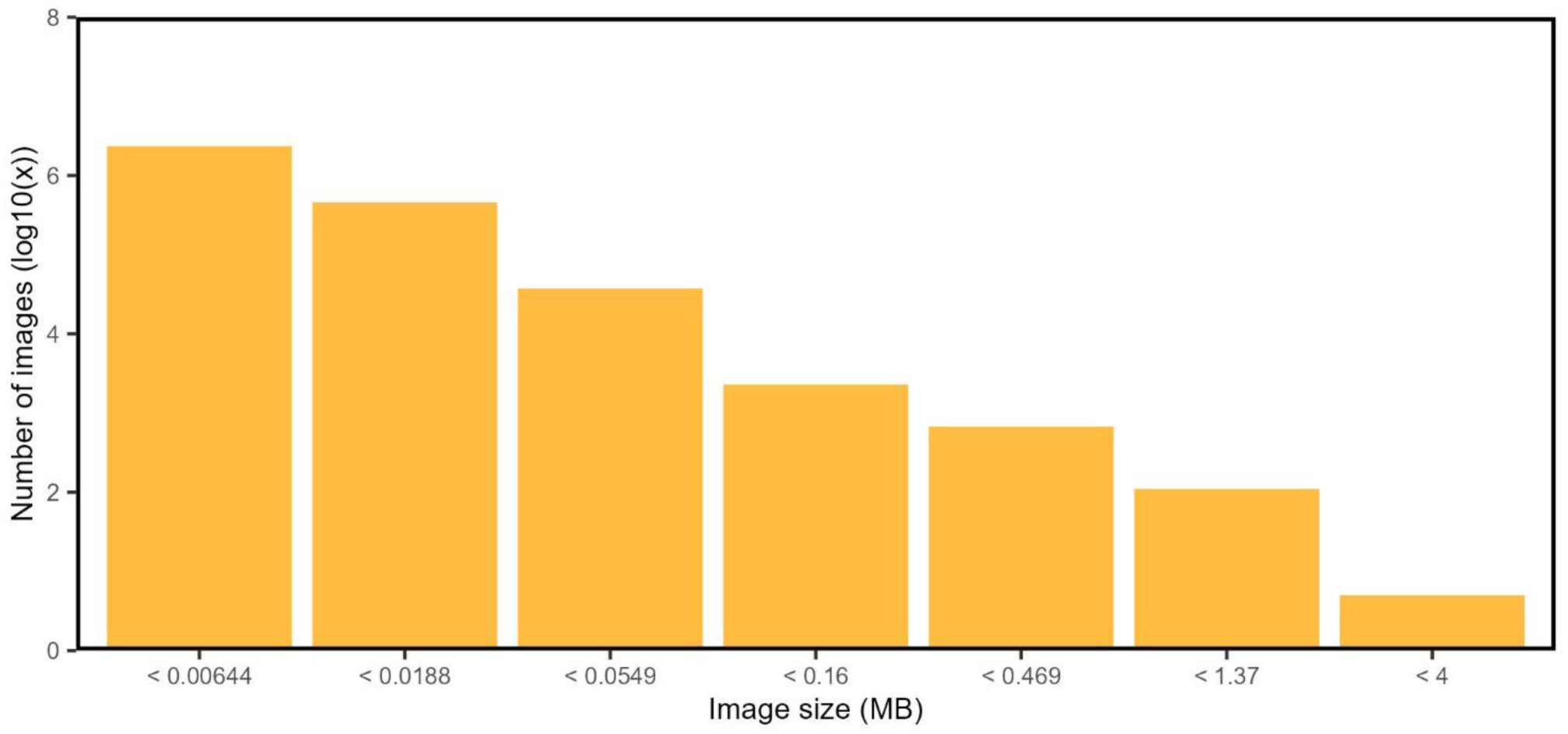
Number of images by storage size (MB), with image size projected onto a logarithmic scale due to skewed distribution towards very small images. The bins include all images less than the size indicated on the y axis but greater than the previous size.

## 5 Discussion

We deployed the Pi-10 continuously over an 18 days period of time. During this time, the instrument was paused briefly twice to allow for cleaning of the flow cell, resulting in an uptime of over 99%.

Our results show that the Pi-10 successfully captures and saves up 5,000 images per minute, when used continuously. When particle density exceeded 147 L^-1^, the instrument stopped saving all images, while still recording the number of particles that passed through the system (i.e. “misses”). We worked mostly in offshore areas where plankton are not the most densely distributed; but were the Pi-10 to be deployed in estuarine or more turbid waters in general, the number of misses would increase with increasing particle density.

The Pi-10 software has a parallel architecture, and in tests with synthetic data can detect 2 million particles per second. In theory, we could therefore capture 100% of the images. This would however result in more than 1 TB an hour. Saving that many images per second was not even attempted, as it is far beyond the capabilities of any reasonable hardware budget. Therefore, one has to make a compromise between storage and data loss. The number of misses in dense concentrations of plankton, is controlled by the user via the “number of saved images per min” setting (see section 2.2.1. Image acquisition software). With the setup deployed here, this selection needs to be based mainly on the speed and size of the disk storage, and to a lesser extent, the speed and capacity of the ship intranet. Instead of sending images across a local network, it is possible to connect a dedicated storage and processing system to the acquisition computer.

However, it’s worth noting that, as per the results, the periods of high misses account for a very small percentage of the data. Furthermore, during periods of high misses / high density we still do record the number of particles passing through the systems, but not the actual images. One can then use the ratio of hits and misses to get a scaling factor. The images that are captured can then be resolved to true numbers by using the scaling factor. The use of a scaling factor in light microscopy is commonplace, with high density samples often being split into 1/512^th^. We very rarely get scaling factors over 252. Thus, the data in these areas is still ecologically usable. As the instrument did not show any bias towards the size of the images that were saved (i.e. it did not prioritise saving smaller images over larger ones or vice-versa), this is akin to subsampling, with more sub-sampling required in areas of high particle density. Sub-sampling is a routine process applied to the analysis of most physical samples collected and it is known that the manual identification process of sub-sampling can over-estimate abundances (Longhurst and Seibert, 1967). One major advantage of using an instrument such as the Pi-10 is that sub-sampling, if not eliminated, is substantially reduced, in turn decreasing potential sub-sampling induced error.

The choice of the range of particle sizes to be collected and saved by the system will also affect the performance of the system and the resulting amount of subsampling needed. In agreement with the documented higher densities of smaller particles (Checkley Jr et al., 2008; Kiko et al., 2022; Stemmann and Boss, 2012), we found the distribution of particle sizes to be highly skewed towards smaller particles. Further increasing the size range of particles collected, by lowering the minimum size (from the 180 µm used here), would result in an increased number of particles captured by the system, beyond the 5000 images per minute capacity. This would translate into more unsaved images and more sub-sampling needed. Furthermore, we’ve found that lowering the minimum size below 100 µm results in more particles than the system can account for, and it crashes as a result. The amount of storage required also increases exponentially with decreasing minimum size.

At a maximum rate of 5,000 images per minute, the digital storage requirements are 32 MB per minute, or 46 GB per day, assuming an average image size of 0.0064 MB as obtained in this study. Therefore, storage space for images collected by the instrument is an important consideration prior to using the instrument. The storage space required will also be dependent on the location of deployment as large images take up more space than small images. In effect, more disk space will be required when the instrument is deployed in areas of high densities of large plankton. The use of portable hard drive may provide an adequate solution, but we found that these are prone to failure when used at sea, possibly due to the vibrations on the ship. Recently, we started using Azure™ disk packs to save the collected images locally prior to uploading them onto cloud storage post survey; this has proven to be a faster and more reliable solution.

It is unquestionable that no plankton sampler can provide a true estimate of abundance for all components of the plankton at any given time (Owens et al., 2013). This is in great part because zooplankton cover such a wide range of sizes, in addition to shapes, and behaviours. As such, each instrument will be biased toward a specific component of the plankton. Whilst the Pi-10 was intended to target the mesozooplankton size range (200 – 20,000 µm), the capacity of the camera system, with a resolution of 10 µm, goes beyond that but is limited by computer processing speed and memory. In the future, as technology continues improving, it will likely be possible for the system to save more than 5000 images per minute. However, we currently recommend using the Pi-10 for what it was intended for, as we have always done successfully with previous versions of the instrument in several studies (Pitois et al., 2018, 2021; Scott et al., 2021, 2023).

It is clear that the number of images generated by the Pi-10 is too many for a human to classify and count manually, without any further subsampling. In earlier studies (Pitois et al. 2018, 2021; Scott et al. 2021), images were extracted at location, subsampled to a manageable level before being classified by an expert taxonomist; essentially replicating a process used with the traditional collection of physical samples followed by microscope analysis. As a result, we could only process a small fraction of the collected dataset. Machine learning tools have been developed to automate this process (Al-Barazanchi et al., 2018; Ellen et al., 2019; Irisson et al., 2022; Luo et al., 2018; Orenstein et al., 2022), many of them are freely available. Using such a tool on the images collected by the Pi-10 is therefore necessary to be able to make the most of the instrument capabilities. While it is possible to embed a classifier within the Pi-10 as part of the instrument, the decision not to do so was taken because: Firstly classification models evolve faster than the hardware on which they are deployed, rendering them obsolete very quickly; Secondly, different users will have different data needs, just like physical plankton samples can be analysed at various taxonomic levels depending on the user’s needs. However, whether these automated tools can achieve a taxonomic resolution that scientists trust, is currently unclear (Giering et al., 2022). So far, we have implemented a classifier first developed by a data study group hosted by the Alan Turing Institute (Data Study Group Team, 2022). This classified images into three classes: copepod, non-copepod and detritus. Copepods were selected as the focus of the classifier due to their prevalence within zooplankton communities, and their importance as food for higher trophic levels. Compromising on taxonomic resolution by selecting only three classes allowed us to build a performant and reliable classifier that is effective to answer our scientific questions. Moving forwards, classifiers will inevitably become more performant and deployed using Edge-AI rather than post-survey. This has been implemented as an example of a UDP stream analyser in a compact Nvidia Jetson computer. The results of this additional study are being prepared for submission in a detailed classifier performance paper.

The main advantage of the Pi-10 is that it can collect high volumes of data in a continuous manner, thus providing unprecedented fine spatial data. This allows us to sample at a much finer resolution than previously possible across larger spatial scales (Scott et al., 2023); thus, opening the door to new areas of research related to the study of small-scale processes which cannot be seen using traditionally collected data, such as plankton patchiness (Plonus et al., 2021) and prey-predator interactions (Suraci et al., 2022). But the pinnacle of pelagic studies will be the monitoring of plankton (phytoplankton and zooplankton) along environmental (e.g., temperature, salinity and other biological (e.g., fish) variables, at high-frequency and in real-time (e.g. Schmid et al., 2023). This offers the opportunity to quickly respond according to observed changes and adjust data collection strategy for example (during a survey for example), but also increases capabilities and potential towards building a digital twin of the oceans (Borja et al., 2024; Chen et al., 2023). Whilst we have been using the Pi-10 on a research vessel, the instrument can be used from any platform connected to water and power supply; thus, offering the opportunity to collect high frequency zooplankton data under different conditions.

## 6 Conclusion

We tested the Plankton Imager Pi-10, an inflow automated mesozooplankton imaging system, aboard the RV Cefas Endeavour and found that it can collect and successfully save up to 5,000 images per minute. At this maximum output, 46 GB of disk space per day are needed to save the collected images assuming an average image size. The high frequency nature of the instrument opens the door to new areas or research, in particular the observation of fine scale processes and the move towards real-time sampling and the increased capability to build a digital twin of the oceans. As technologies, including hardware (e.g. camera optics, computer chips) and software (algorithms based on computer vision) continue evolving and become more performant, the Pi-10 performance will also improve, being able to collect and save more and more images that will be confidently classified to a high taxonomic resolution.

## Acknowledgments

We acknowledge the considerable efforts of scientists, officers and crew of RV Cefas Endeavour involved in all surveys where the Plankton Imager was deployed.

